# What Determine the Distribution and Richness of Wild Spices in the Sudano-Guinean zone of Benin, West Africa?

**DOI:** 10.1101/680660

**Authors:** K.M. Kafoutchoni, R. Idohou, K.V. Salako, C. Agbangla, A.E. Assogbadjo

## Abstract

In Benin, most of the spices used for food, medicine and ceremony are gathered from the wild as little attempt has been made so far for their domestication and cultivation. Consequently, many wild spices are prone to overexploitation and threatened by habitat loss. Also, little information is available regarding their occurrence areas and the factor determining their geographical distribution and richness. This study aimed at i) mapping the distribution and the richness of 14 wild spices, and ii) assessing the drivers of their distribution and richness patterns in the Sudano-Guinean zone of Benin. Data were collected during field exploration and from the database of the Global Biodiversity Information Facility. Species distribution was mapped, and a grid of 10 × 10 km cells and a circular neighborhood option with a radius of 10 km was used to assign points to grid cells, then species richness was mapped. The species were unequally distributed across the study area. High species richness occurs in Bassila and Zou phytodistricts. Three spice-rich areas are needed to capture all the wild spices at once. Interaction of mean temperature of driest quarter, altitude, and precipitation seasonality significantly shaped the distributional range of three wild spices (*Aframomum alboviolaceum, Uvaria chamae* and *Zanthoxylum zanthoxyloides*), while the same factors in addition to clay content between 5-15 cm, contributed significantly to create appropriate conditions for the cooccurrence of several species.

**Résumé:** Au Bénin, la plupart des épices utilisées pour l’alimentation, en médecine et pour les cérémonies sont collectées dans la nature car très peu de tentatives ont été faites pour leur domestication et leur culture. Par conséquent, de nombreuses espèces d’épices sauvages sont surexploitées et se retrouvent menacées par la destruction de leur habitat. Aussi, peu d’informations sont disponibles sur leurs zones d’occurrence et les facteurs influençant leur distribution géographique et leur richesse. Cette étude visait à i) cartographier la distribution et la richesse de 14 épices sauvages, et à ii) évaluer les facteurs influençant leurs distribution et richesse dans la zone Soudano-Guinéenne du Bénin. Les données ont été collectées au cours d’explorations sur le terrain et à partir de la base de données du Global Biodiversity Information Facility. La distribution des espèces a été cartographiée et une grille de 10×10 km a servi de base pour la cartographie de la richesse en espèces. Les espèces étaient inégalement distribuées dans la zone d’étude. Des zones de grande richesse en épices sont présentes dans les phytodistricts de Bassila et du Zou. Trois zones de forte diversité en épices sont nécessaires pour capturer toute la diversité du groupe fonctionnel des épices sauvages. L’interaction de la température moyenne du trimestre le plus sèche, l’altitude, et la saisonnalité des précipitations ont significativement influencé la distribution de trois épices sauvages (*Aframomum alboviolaceum, Uvaria chamae* et *Zanthoxylum zanthoxyloides*). Ces trois facteurs, ajoutés au taux d’argile dans le sol, ont contribué à la création des conditions favorables pour la cooccurrence de plusieurs espèces d’épices sauvages.

**Mots clés:** arbre d’inférence conditionnelle, épices sauvages, espèces négligées et sousutilisée, SIG, systèmes d’information géographiques,

## 1. Introduction

In recent decades, increasing attention has been given to Non-timber forest products (NTFPs) due to the key role they play in the livelihood of rural communities. Indeed, NTFPs ensure food security, provide medicine and cash income to poor rural people, mainly in periods of drought and starvation (Sanchez *et al*., 2012). So far, studies on NTFPs especially wild plant species have known a huge boost. In Africa, some of such works relate to the functional group of wild spices and have been conducted mainly in Nigeria (Belewu *et al*., 2009) and Cameroon (Etoundi *et al*., 2010). Nonetheless, there still exists in the nature many wild plants including spices, not yet scientifically studied despite their great potential. In Benin, many spices used for medicine, food and ceremony are gathered from the wild. Most of these wild spices are reported to be effective in the treatment of various ailments such as fever, urinary disorder, snakebite, hypertension, toothache, menstrual disorder, and diabetes (Srinivasan, 2005). Moreover, several wild spices contain key mineral salts including phosphorus, iron, calcium and magnesium, and can thus be used to tackle malnutrition among rural communities (Al-Jasass & Al-Jasser, 2012). Wild spices are also rich in different bioactive compounds (alkaloids, polyphenols, flavonoids, steroids, essential oils, etc.) that give them their medicinal properties (Belewu *et al*., 2009; Etoundi *et al*., 2010). In spite of their importance, little attempt has been made so far for the domestication and cultivation of wild spices. Consequently, many spices are being extinct due to overexploitation and habitat loss. Moreover, very little is known about the geographical distribution of the wild spice resources and how local ecological conditions shape their distribution and richness patterns in Benin. However, conservation and sustainable management of these resources require to understand how the species are distributed as well as the factors determining their distribution and richness (Wallace, 2011), in order to develop appropriate strategies for each species. Geographic information systems (GIS) has emerged as a powerful tool to help ecologists and conservationists identify areas of high diversity, and has been successfully used to determine high diversity areas in wild potatoes (Hijmans & Spooner, 2001), piper (Parthasarathy *et al*., 2006), and Asteraceae in Iran (Zare *et al*., 2013). This study aimed to i) map the distribution and the richness of 14 wild spices, and ii) assess the drivers of their distribution and richness patterns in the Sudano-Guinean zone of Benin.

## 2. Methodology

### 2.1. Study area

The study was conducted in the Sudano-Guinean zone of Benin located between the latitudes 7°30’ and 9°45’ N. The region is subdivided into three phytodistricts (Bassila, Zou and South-Borgou) characterized by different vegetation and environmental conditions (Adomou *et al*., 2006). The mean annual rainfall varies between 1100 mm and 1300 mm. The annual temperature varies from 25°C to 29°C and the relative humidity varies between 31% and 98%. The soils are ferruginous with variable fertility. The vegetation consists in a mosaic of woodland, dry dense forests, gallery forests, as well as tree and shrub savannas.

### 2.2. Data collection

The data used in this study are part of an ongoing research on the diversity and ethnobotany of the wild spices in Benin. The overall methodology included an exploratory survey to identify localities of effective presence of wild spices, random selection of 10 localities across the three phytodistricts for surveys, individual interviews and field visits. These data were collected on 14 wild spices inventoried during field exploration in 2016. The species include *Aeollanthus pubescens*, *Aframomum alboviolaceum*, *Aframomum angustifolium*, *Aframomum melegueta*, *Clausena anisata*, *Cymbopogon giganteus*, *Lippia chevalieri*, *Monodora tenuifolia*, *Piper guineense*, *Securidaca longipedunculata*, *Thonningia sanguinea*, *Uvaria chamae*, *Xylopia aethiopica*, and *Zanthoxylum zanthoxyloides*. Occurrence of each species were recorded using a GARMIN GPSMAP 62s. Additional occurrence data were retrieved from the Global Biodiversity Information Facility (https://www.gbif.org). Environmental data included the full set of 19 bioclimatic variables downloaded at 1km resolution from <u>Worldclim</u>, three layers of six soil characteristics at 250 m resolution obtained from the Africa Soil Profiles Database (<u>ISRIC</u>), and elevations. Soil characteristics considered were soil organic carbon (g/kg), pH in H2O, clay content (%), sand content (%), silt content (%), cation exchange capacity (cmol/kg) and bulk density (tons/m^3^), each at 0-5 cm, 5-15 cm and 15-30 cm of depth.

### 2.3. Data analysis

Occurrence data were used for mapping the distribution of each species. The study area was subdivided into 618 cells of 0.1 degree^2^ resolution and in each cell the number of species was computed. To generate an accurate richness map, less biased by the subjective assignment of grid origin and less sensitive to small errors in the geographical coordinates, species were assigned to grid cells using a circular neighborhood (Ocampo *et al*., 2010) of radius 10 km. A Conditional inference tree (CIT) model was used to determine which ecological factors drive the most the distribution and the richness patterns of the wild spices, and how. Tree-based models such as CIT are reliable in examining patterns in ecological data, particularly when hierarchical interactions exist between the factors (Johnstone *et al*., 2014). As the respective spatial resolutions of climate and soil data were different, bioclimatic variables were downscaled to the resolution of 250 m using the bilinear interpolation algorithm in ArcMap 10.3. To deal with common inter-collinearity in ecological data (Zuur *et al*., 2010), bivariate Pearson’s correlation coefficients were computed for each dataset and highly correlated variables (r>0.8) were grouped. Then the most biologically meaningful variable was selected from each group. The conditional inference tree was generated with the package *partykit* (Hothorn & Zeileis, 2005) in R. All maps were generated using ArcMap.

## 3. Results and discussion

### 3.1. Geographical distribution and species richness map

The distribution map of the wild spices showed an uneven geographical distribution throughout the Sudano-Guinean zone (Figure 1). *Aeollanthus pubescens, Aframomum alboviolaceum, Cymbopogon giganteus, Lippia chevalieri, Monodora tenuifolia, Uvaria chamae* and *Zanthoxylum zanthoxyloides* were widely distributed across the three phytodistricts, although the distribution of *L. chevalieri* followed an aggregative pattern. *Aframomum angustifolium* and *Thonningia sanguinea* occurred everywhere but the geographical repartition of the first was relatively less extensive while the second occurred mostly in Bassila and Zou phytodistricts. *Clausena anisata* and *Xylopia aethiopica* were found both in Zou and Bassila phytodistricts except that occurrence of *X. aethiopica* was lesser. *Securidaca longipedunculata* was well distributed in Borgou-South and Zou phytodistricts whereas *Aframomum melegueta* and *Piper guineense* were rare and found exclusively in the sacred grove of Badjamè in the extreme Southwest of the Zou phytodistrict (Fig. 1).

**Fig. 1.**
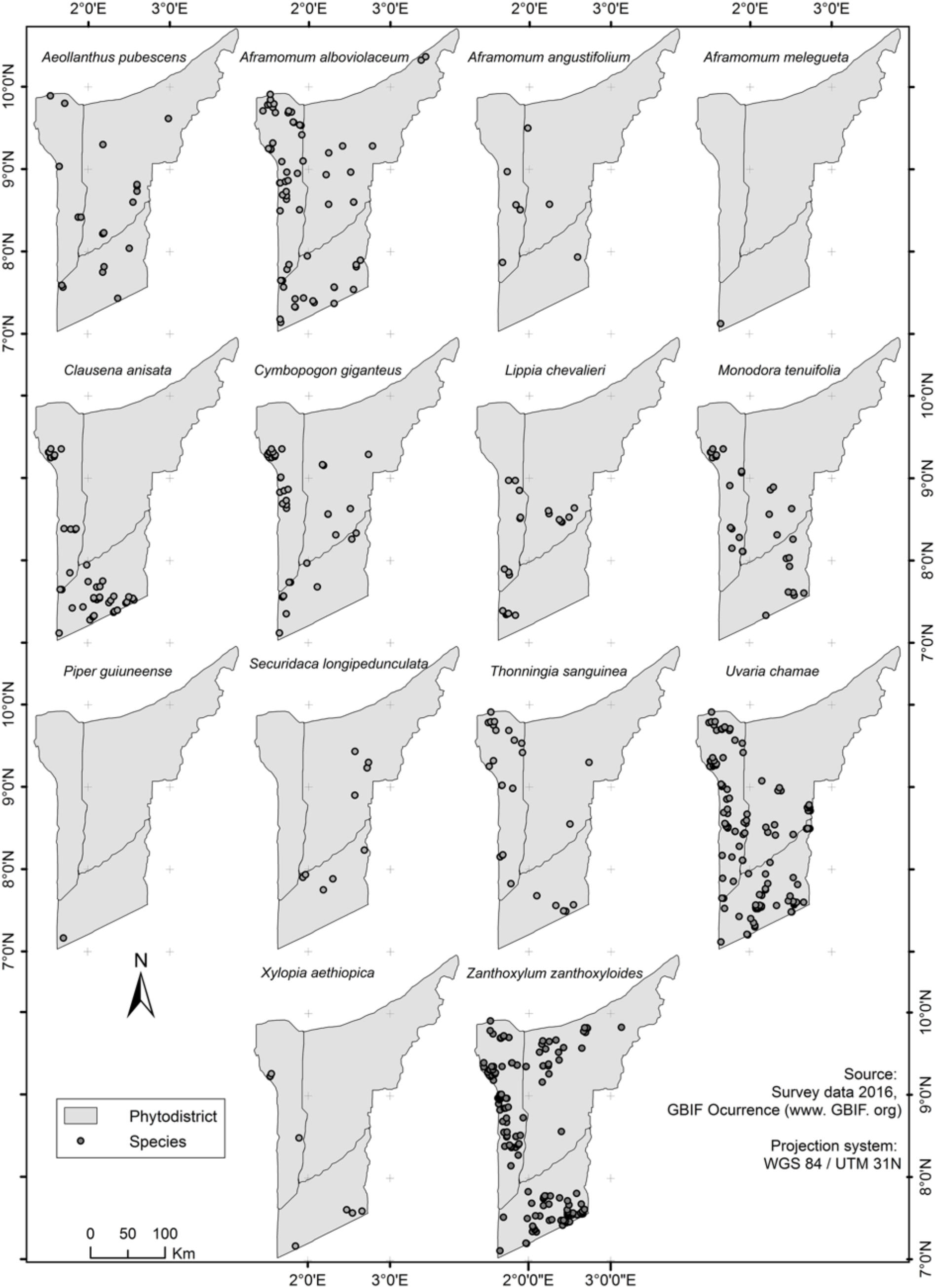
Map showing the distribution of 14 wild spices found in the Sudano-Guinean zone of Benin

Species richness was not homogeneously distributed across the phytodistricts (Fig. 2). Southwards, the first spice-rich area was located in Bassila and extended between 1°26’40”-1°48’29”E and 8°50’43”-9°24’39”N, with 11 species (*A. pubescens, A. alboviolaceum, A. angustifolium, C. anisata, C. giganteus, L. chevalieri, M. tenuifolia, T. sanguinea, U. chamae, X. aethiopica* and *Z. zanthoxyloides*). The second area, located between 1°44’36”-2°1’7”E and 8°21’15”-8°31’0”N, overlapped Bassila and South-Borgou phytodistricts and included 10 species namely *A. pubescens, A. alboviolaceum, A. angustifolium, C. anisata, C. giganteus, L. chevalieri, M. tenuifolia, S. longipedunculata, U. chamae* and *Z. zanthoxyloides*. Area 3 with lesser extent, was localized in South-Borgou and consisted in 8 species (*A. alboviolaceum, A. angustifolium, C. giganteus, L. chevalieri, M. tenuifolia, T. sanguinea, U. chamae* and *Z. zanthoxyloides*). Area 4 was a small extent overlapping Bassila and Zou, and consisting in 8 wild spices (*A. alboviolaceum, C. anisata, C. giganteus, L. chevalieri, S. longipedunculata, T. sanguinea, U. chamae* and *Z. zanthoxyloides*). The fifth area was a stretch between 2°2’37”-2°31’8”E and 7°26’27”-7°48’58”N with 10 species namely *A. pubescens, A. alboviolaceum, C. anisata, C. giganteus, M. tenuifolia, S. longipedunculata, T. sanguinea, U. chamae, X. aethiopica* and *Z. zanthoxyloides*. The spice-rich area was located in southwestern of Zou and contained 8 species including *A. melegueta* and *P. guineense*, two wild spices exclusively encountered in this phytodistrict. Although the first area was enough to capture about 79% (11 species) of the wild spices found in the Sudano-Guinean zone, three spice-rich areas including areas 1 and 6 will be required to conserve the whole diversity of the functional group.

**Fig. 2.**
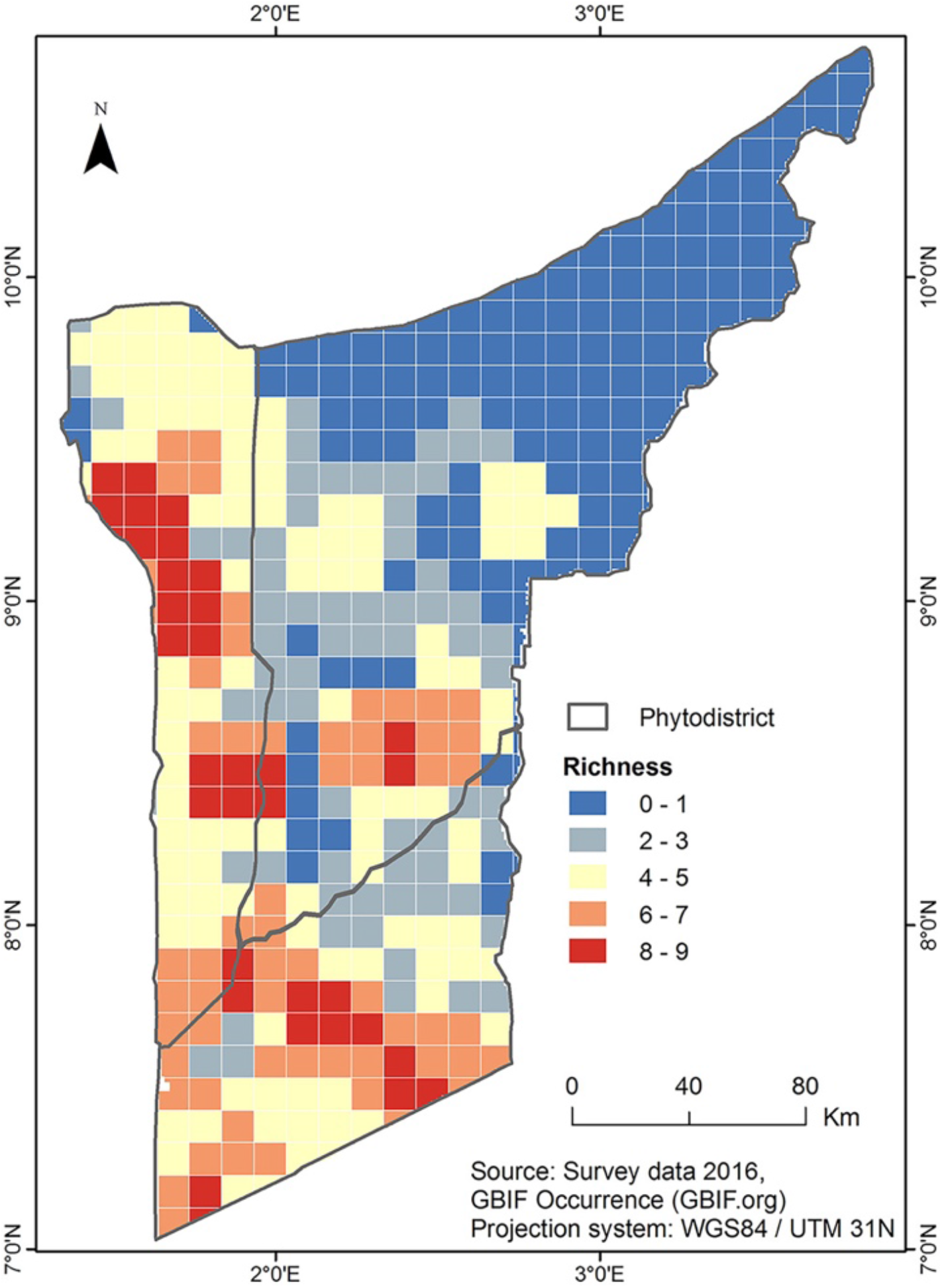
Richness map of 14 wild spices in the Sudano-Guinean Zone of Benin

To some extent the spices richness map generated in this study was consistent with conclusions from a previous study of Adomou *et al*. (2010). Based on overall species richness, threatened species concentration, presence of endemic plant species and threatened plant communities, the authors worked out several areas as high priority vegetation sites for plant conservation in Benin. These sites included the protected forest of Pénéssoulou, Bassila and Monts Kouffé and the unprotected site of Djagbalo, which correspond to the two spice-rich areas identified in the Bassila phytodistrict.

### 3.2. Assessing the factors determining wild spices distribution and richness patterns

Four bioclimatic variables and six soil characteristics were less correlated (r < 0.8) and used together with altitudes to generate the Conditional inference tree (CIT). Selected bioclimatic variables were maximum temperature of warmest month (BIO 5), mean temperature of driest quarter (BIO 9), precipitation of driest month (BIO 14) and precipitation seasonality (BIO 15). Less correlated soil characteristics were cation exchange capacity (cec 2), clay content (clay 2), silt content (silt 2), pH in H2O (ph2), all between 5-15 cm depth, and organic carbon at 0-5 cm (oc1) and 5-15 cm (oc2). CIT model showed that among these explanatory variables, BIO 9, BIO 15, clay2, oc1 and altitude were important in explaining the distribution and richness patterns of the species (Fig. 3). Mean temperature of driest quarter (BIO 9) highly significantly (p < 0.001) contributed in splitting the tree at two consecutive levels (nodes 1 and 2) with threshold values of 28.2° C and 27° C respectively. For BIO 9 values less or equal to 27° C, altitude was significant (p = 0.011) in differentiating the species at the threshold of 299 meters (node 3). Altitudes ≤ 299 m were very suitable for the distribution of *U. chamae* and to a little extent for that of *Z. zanthoxyloides* (node 4), while above 299 m *Z. zanthoxyloides* had a relatively good distribution (nodes 6 and 7). For mean temperature of driest quarter between 27° C - 28.2° C, conditions were fairly (p = 0.05) advantageous for many species, particularly for *A. alboviolaceum* when variation in precipitations (BIO 15) was lesser or equal to 59%, and for *U. chamae* and *Z. zanthoxyloides* when BIO5 > 59% (nodes 9 and 10). For BIO 9 values greater than 28.2° C, *Z. zanthoxyloides* thrived exceptionally better at altitudes bellow or equal to 66 m (nodes 13 and 14) whereas conditions were suitable for the occurrence of *U. chamae* when the altitude is upper than 66m (node 15; Fig. 3). Therefore, interaction of mean temperature of driest quarter, altitude, and precipitation seasonality determine the distributional range of three wild spices over the fourteen under study. This finding is in agreement with the general statement that the distribution of wild plant is shaped by the interaction of different ecological factors (Šímová *et al*., 2011).

**Fig. 3.**
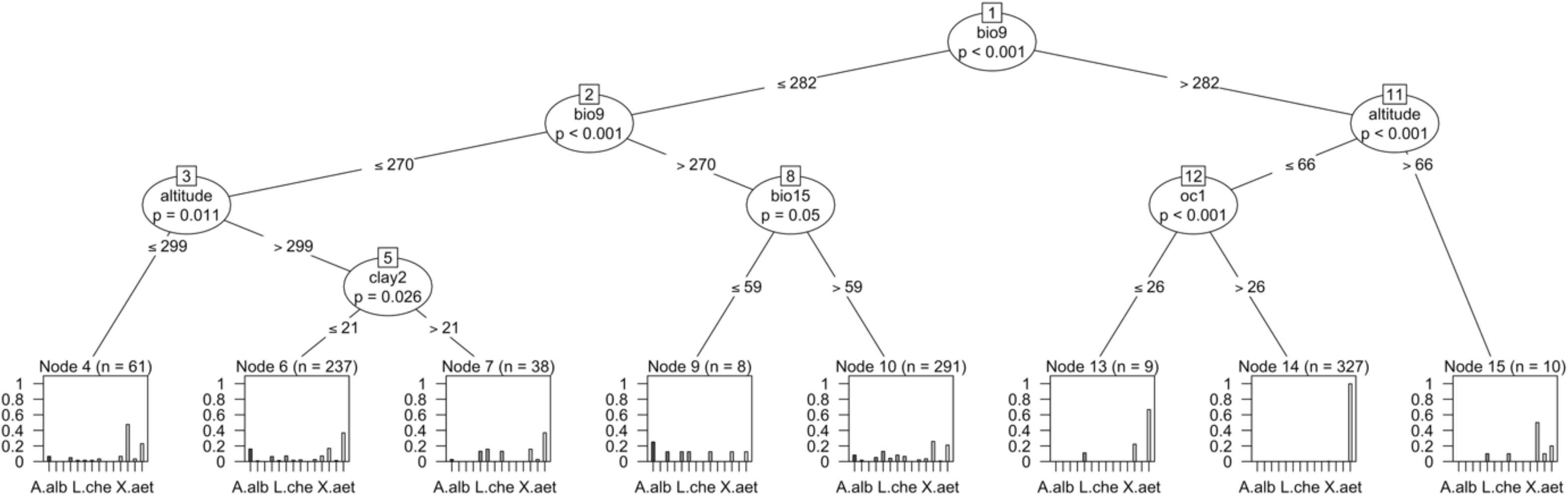
Conditional inference tree result for wild spices distribution and richness patterns. Bars represent the proportion of species in each particular ecological condition

Based on the results, appropriate zones can be selected for the cultivation or domestication of these species. For instance, *Z. zanthoxyloides* a widely consumed wild, can be cultivated almost everywhere in the study area since combination of the mean temperature of driest quarter, altitude and precipitation seasonality across the phytodistricts met its preferences. This fully supports the ecology of the species whose origin is Sudano-Guinean and which is thus found in all three phytodistricts of the study area. In the same way, our results suggested that *U. chamae* can be preferentially cultivated in Zou phytodistrict and up to the latitude of Bantè (8°24’N), whereas *A. alboviolaceum* is shown to perform well within a stretch in extreme Southwestern Zou. Although *U. chamae* is a Guineo-Congolian species, it has a wide geographical distribution in Benin, even extending to the far Northern. The species is thus adapted to a large climatic gradient and thereof the identified ecological determinants of its distribution may not actually affect the aforementioned distribution enough. On the other hand, the wide distribution of *U. chamae* across the Sudano-Guinean zone, out of its ecological range can be due to human dissemination as well, especially in sociolinguistic areas of Idatcha, as these people consume the root barks of the species as vegetable in soup (Achigan-Dako *et al*., 2010). Being an afrotropical species, *A. alboviolaceum* has large ecological and geographical amplitudes and is then less sensible to temperature and precipitation, as most tropical regions experience little intra-annual climatic variability (Blach-Overgaard *et al*., 2010). However, the species thrives better in savannas, explaining its wide occurrence throughout the Sudano-Guinean zone.

Overall, conditional inference tree revealed that altitude, precipitation seasonality (BIO 15) and clay content at 5-15 cm (clay 2), each along with mean temperature of the driest quarter (BIO 9), contributed significantly in creating appropriate environmental conditions for the cooccurrence of several species to create spice-rich areas. Thus, up to ten wild spices can potentially flourish in areas with a mean temperature of driest quarter BIO 9 ≤ 27 ° C and where altitude was lower or equal to 299 m. Similarly, seven to twelve species thrived more or less well in areas where mean temperature of driest quarter and altitude were respectively greater than 27° C and 299 m, and depending on the clay content in soil at 5-15 cm which threshold was 21% (nodes 6 and 7; Fig. 3). Additionally, coefficient of variation of precipitation (BIO 15) significantly (p = 0.05) contributed to the establishment of spice-rich areas harboring from seven to twelve species at 59% of threshold, when mean temperature of driest quarter was higher than 27° C (nodes 9 and 10; Fig. 3).

## 4. Conclusion

The study gives insight into the driving forces of the distribution and richness patterns of 14 wild spices in Benin. It appeared that the interaction of mean temperature of driest quarter, altitude, and precipitation seasonality significantly shape the distributional range of three wild spices (*Aframomum alboviolaceum, Uvaria chamae* and *Zanthoxylum zanthoxyloides*). Regarding species richness, clay content between 5-15 cm, in addition to these three variables, contributed significantly to create appropriate environmental conditions for the co-occurrence of several species to create spice-rich areas.

